# Discovery of cyclic peptide natural product inhibitors of *Balamuthia mandrillaris*

**DOI:** 10.1101/2024.05.03.592372

**Authors:** Chenyang Lu, Samantha Nelson, Gabriela Coy, Christopher Neumann, Elizabeth I. Parkinson, Christopher A. Rice

**Author notes:** These authors contributed equally to this work (Chenyang Lu, Samantha Nelson).

## Abstract

*Balamuthia mandrillaris* is a pathogenic free-living amoeba that causes infection of central nervous system, called *Balamuthia* amoebic encephalitis (BAE), as well as cutaneous and systemic diseases. Patients infected with *B. mandrillaris* have a high mortality rate due to the lack of effective treatments. A combination of non-optimized antimicrobial drug regimen is typically recommended; however, they have poor parasite activity and can cause various severe side effects. Cyclic peptides exhibit a broad spectrum of antimicrobial activities and lower cytotoxicity. In this study, we evaluated the anti-*B. mandrillaris* effect of cyclic peptides. The predicted natural product-43 (pNP-43), identified from the SNaPP (Synthetic Natural Product Inspired Cyclic Peptides) library, and its derivates displayed a significant inhibition for *B. mandrillaris* trophozoites. Eight pNPs had IC_50_s <5 μM. Furthermore, all hit pNPs demonstrated minimal hemolytic and cytotoxic effects on human cells. Our study first indicates the anti-*B. mandrillaris* effect of cyclic peptides, which provides a new direction for drug development. Further studies of the mechanism of action and *in vivo* effects will be elucidated to confirm the potency as a treatment for *B. mandrillaris* infection in the future.

## Introduction

*Balamuthia mandrillaris* is a pathogenic and opportunistic free-living amoeba (FLA), which causes infections in humans and animals. It can initially cause cutaneous skin lesions followed by infection of central nervous system (CNS) called *Balamuthia* amoebic encephalitis (BAE). However, most cases in the US only report CNS symptoms without skin lesions, which develop quicker than when cutaneous disease presents. Time to death from neurological complications varies from weeks to months. In the United States, 109 cases were reported from 1974-2016, with the majority of cases being concentrated in the southwestern regions, with 55% of these cases being within the Hispanic population.^1^ Another hot spot for BAE is Peru, which has had 55 cases reported since 1975.^2^ With the increased awareness, knowledge of at-risk populations and development of more specific diagnostic tools (next-generation sequencing), more cases have been identified in the past 20 years, with 28 cases reported in China,^3^ as well as cases in India, Australia, Europe, East Asia and Africa. ^4–8^

For BAE, there are no FDA-approved therapies. When patients are diagnosed, they have been administered with various therapeutic regimens, often using a combination of antimicrobial drugs and drugs that were used in previously successful cases (**Figure 1**). The CDC recommended treatment consists of flucytosine, pentamidine, fluconazole, sulfadiazine, azithromycin, clarithromycin and/or miltefosine. However, the existing therapeutics are still unsuccessful, with the mortality rate in the United States being ∼92%.^1^ Overall, this high incidence of drug failure points to the great need for effective anti-*B. mandrillaris* therapeutics. While a few drug screens have been performed to identify novel compounds, these screens have been primarily libraries of small molecules or specific natural products such as curcumin or resveratrol.^9–15^

**Figure 1.**
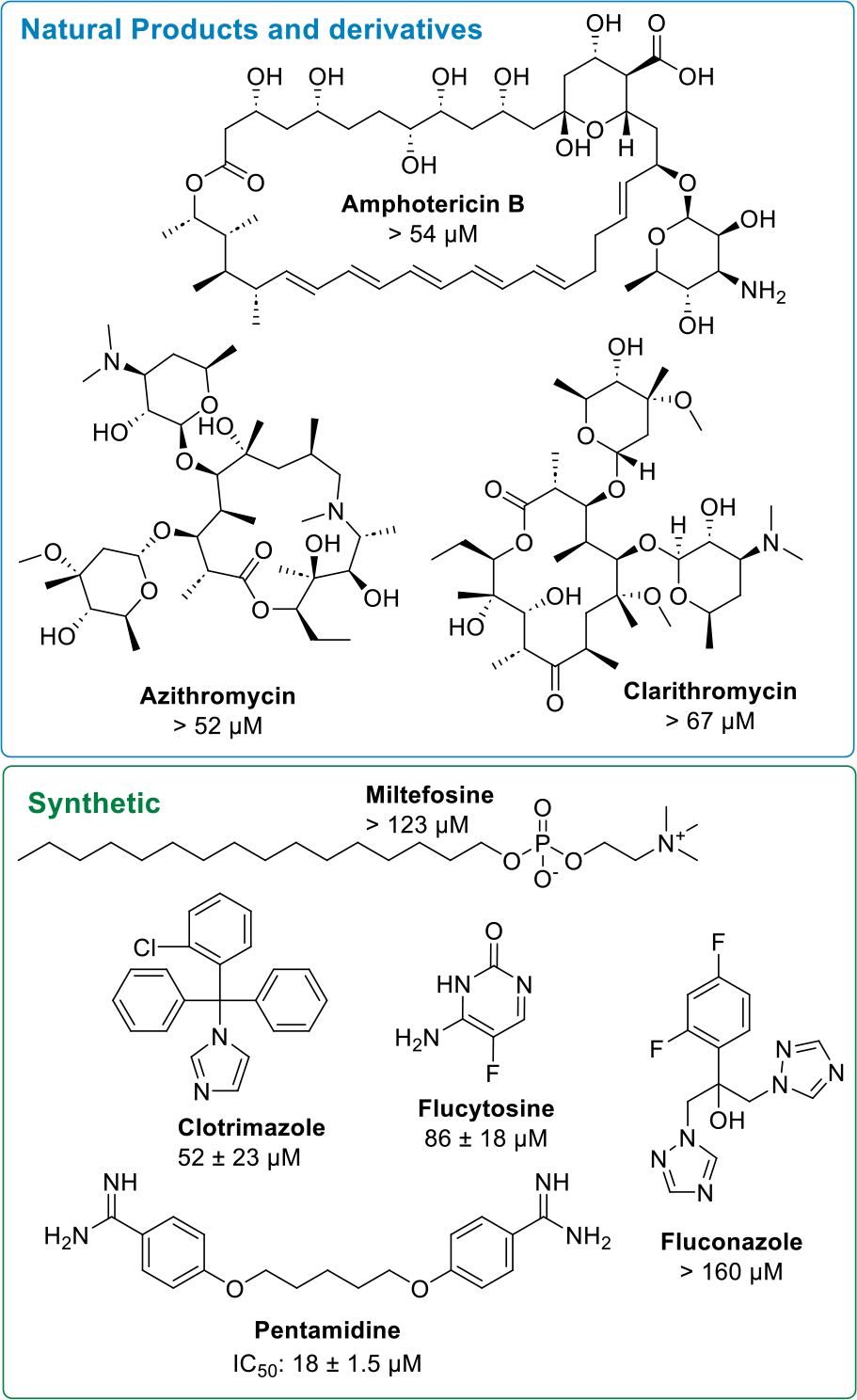
Structures and IC_50_s of CDC recommended treatments for *B. mandrillaris* infections.

Cyclic peptides are a growing class of therapeutics with 18 approved by the FDA in the past two decades.^16^ Cyclic peptides are a promising source of a therapeutics because they reside in a unique space that balances the characteristics of both small molecules and biologics, known as the Goldilocks space.^17^ They are capable of hitting challenging targets such as protein-protein interactions and often have similar specificity to biologics while having bioavailabilities and cell permeabilities more similar to small molecules^17,18^. Furthermore, cyclic peptides have increased cell permeability and proteolytic stability compared to their linear counterparts^19,20^.Peptides have demonstrated activity against multiple disease states including, antifungal, antibacterial, immunosuppressants, anticancer, and more.^16,21,22^ Some peptides have also been shown to have activity against FLAs (i.e. *N. fowleri, A. globifora*, and *A. castellanii*),^23– 28^ but none have demonstrated activity against *B. mandrillaris*. Specifically, a tyrocidine-derived linear peptide exhibited IC_50_ against *A. castellanii* of 112 μg/mL (88 μM)and 82 μg/mL (64 μM) against *N. fowleri* at 24 h.^24^ Nisin, a ribosomally synthesized post translationally modified peptide produced by *Lactococcus lactis*, appears to affect the growth of *Acanthamoeba*, resulting in smaller and rounder trophozoites. However, the trophozoites proliferated again after 72 h. Overall, the IC_50_ of nisin was quite poor (4493.2 IU/mL, 4.49 mg/mL at 24 h)^28^ suggesting that it is unlikely to be an effective treatment for *Acanthamoeba*.

Given that little is known about the activity of cyclic peptides against *B. mandrillaris*, we decided to explore their potential to treat this challenging disease. Previously, we developed a library of natural product-like cyclic peptides known as synthetic natural product inspired cyclic peptides (SNaPP).^29^ The SnaPP library was generated by predicting structures of nonribosomal peptides from cryptic nonribosomal peptide biosynthetic gene clusters that also contain a penicillin binding protein (PBP)-type thioesterase, a class of enzyme only known to catalyze head-to-tail cyclization of nonribosomal peptides. SnaPP is made up of head-to-tail cyclized predicted natural product peptides (pNPs) ranging in size from 4 to 10 amino acids. The SNaPP library has previously been screened for antibiotic activity, resulting in the identification of cyclic peptides with promising activities against Gram-positive and Gram-negative bacteria.^29^ More recently, we have found cell permeable cyclic peptides that stimulate the proteasome^30^. In this paper, we have screened the SNaPP library for activity against *B. mandrillaris* using a previously developed protocol for trophocidal activity.^9^ This has enabled us to identify active cyclic peptides with half maximal inhibitory concentrations (IC_50_s) well below current standard of treatment for *B. mandrillaris* that also display minimal-to-no cytotoxicity to mammalian cells.

## Methods

### Maintenance of *Balamuthia mandrillaris*

The pathogenic *Balamuthia mandrillaris* (CDC V039; ATCC 50209), a BAE strain, was isolated from a pregnant baboon at the San Diego Zoo, San Diego, USA, in 1986. *B. mandrillaris* trophozoites were routinely grown axenically at 37°C in *Balamuthia mandrillaris* ITSON (BMI) medium (bacto casitone, 20 g/liter; Hanks’ balanced salts, 68 mL/liter and 912 mL of deionized water). After autoclaving, 10% fetal bovine serum and 1% penicillin/streptomycin antibiotics were added to produce the complete BMI. All experiments were performed using logarithmic phase trophozoites. Before experimentation, 0.25% Trypsin-EDTA (Gibco, Gaithersburg, MD, USA) was used to detach the cells from the vented 75 cm^2^ tissue culture flasks (Olympus, El Cajon, CA, USA) and the cells were collected by centrifugation at 3214 RCF.

### *In Vitro* CellTiter-Glo Trophocidal Assay

The trophocidal activity of cyclic peptides was assessed using the CellTiter-Glo 2.0 (Promega, Madison, WI) luminescent viability assay as previously described.^9,11^ In brief, all cyclic peptides were diluted in BMI and serially diluted in 2-fold dilutions 6 times with a final concentration of 10% DMSO in clear 96-well plates (Costar 3370) “mother plates”. After serial dilutions were performed, 10 μL of peptides, from the diluted stock plates, were put into white 96-well plates (Thermo Fisher Scientific, #136102, USA) “assay ready plates”. To combine parasites for screening, 90 μL of *B. mandrillaris* trophozoites were seeded at 16,000 cells/well from of a stock concentration of 1.77 x 10^5^/mL for a total time of 72 hours at 37 °C. DMSO (1%) and 90 μM of atorvastatin were used as the negative and positive control, respectively. At the 72-h time point, 25 μL of CellTiter-Glo 2.0 reagent was added to all wells of the white 96-well plates. The plates were shaken at 250 rpm in a dark environment for 2 minutes to induce cell lysis, then incubated at room temperature for 10 min to stabilize the luminescent signal. The adenosine triphosphate (ATP) luminescent signal (relative light units; RLUs) was measured at 490 nm with a SpectraMax I3X plate reader (Molecular Devices, Sunnyvale, CA, USA). Each peptide was tested in a minimum of three biological replicates using different populations of cells. Half maximal inhibitory concentration (IC_50_) curves were generated using total ATP RLUs, where controls were calculated as the average of replicates using the Levenberg–Marquardt algorithm, using DMSO as the normalization control, as defined in CDD Vault (Burlingame, CA, USA).

### Mammalian Cytotoxicity Counter Screening

Cytotoxicity of the cyclic peptides was also determined by using the CellTiter-Glo 2.0 assay on A549 human lung carcinoma cells. A549 cells were seeded at a concentration of 1.6 × 10^4^ cells/mL in 96-well plates (Thermo Fisher Scientific, USA), in the presence of serially diluted compounds, as described above. Again, 1% DMSO, and 100 μM of chlorhexidine were used as the negative and positive controls. A549 were grown in F12K medium (Fisher Scientific, USA) supplemented with 10% FBS and 1% penicillin/streptomycin antibiotics. The peptide concentration was screened from 100 μM to 3.125 μM. The total volume of each well was 100 μL, containing 10 μL compounds and 90 μL cells, and plates were incubated at 37 °C, 5% CO_2_ for 72 h. At the 72-h time point, CellTiter-Glo 2.0 reagent was added to all the wells and read by the SpectraMax I3X plate reader as described above. Curve fitting using non-linear regression was carried out using the average of replicates and the Levenberg–Marquardt algorithm, using DMSO as the normalization control, as defined in CDD Vault. We then calculate the selectivity index (SI) by using the equation; SI = (IC_50_ of A549’s)/(IC_50_ of *B. mandrillaris*) to determine specificity of inhibitors. A drug with a SI value ≥ 10 was considered the minimum standard for further evaluation as a hit drug.

### Statistical Analysis

The Z’ factor was used as a statistical measurement to assess the robustness of each drug plate. This factor uses the mean and standard deviation values of the positive and negative controls to assess data quality. The robustness of all of the plates screened had an excellent Z’-score value of 0.75 or above.

### Hemolysis Assay

In accordance with a previously delineated methodology,^31^ whole human blood was procured from BiolVT and utilized prior to its expiration date. 100 μL of blood was combined with 500 μL of sterile 0.9% NaCl in a 1.5 mL Eppendorf tube, followed by thorough mixing through inversion. The resulting mixture underwent centrifugation at 500 rcf for 7 minutes, after which the supernatant was extracted using a pipette. The pellet was subjected to two additional washes, and following the final wash, 800 μL of Red Blood Cell (RBC) buffer was added to resuspend the red blood cells. Subsequently, 4 μL of the compounds (ranging from 100 to 0.78 μM in DMSO, final concentration), 76 μL RBC buffer, and 40 μL of resuspended red blood cells were added to a 96 U-well plate (VWR) and then incubated for 1 hour at 37°C. The plate underwent centrifugation at 500 g for 5 minutes, after which 75 μL of the supernatant was transferred to a 96 flat-well plate for absorbance measurement at 540 nm using a SpectraMax iD3 plate reader. Percent hemolysis was determined based on the average absorbance of the positive (triton-X 100) and negative controls (DMSO), with a minimum of three biological triplicates conducted.

### Resin Loading

Utilizing a previously outlined approach,^31^ 2-chlorotrityl chloride (2-CTC) resin (1g, with a loading capacity of 0.56 mmol/g, mesh size 100-200), underwent swelling in Dimethylformamide (DMF) for 5 minutes. After the removal of DMF, the swollen 2-CTC resin was exposed to fluorenylmethyloxycarbonyl (Fmoc)-protected amino acid (3 equiv.), N, N-Diisopropylethylamine (DIPEA) (4 equiv.), and DMF (0.07 mM) for a duration of 2 hours. Subsequently, the solution was drained, and a mixture comprising of Dichloromethane (DCM): Methanol (MeOH): DIPEA in a ratio of 17:2:1 (0.07 mM) was employed to treat the resin, followed by agitation for 30 minutes to cap any remaining unreacted resin. After draining the solution, the resin underwent filtration and was washed thrice with DCM (5 mL each) and MeOH (5 mL each) before being dried for 1 hour. Subsequently 1.5-3.0 mg of dried resin was exposed to a 20% mixture of piperidine/DMF (500 μL) for 15 minutes. Further, 100 μL of the resulting supernatant was added thrice to 900 μL of DMF. The reaction mixtures were subsequently analyzed via UV absorbance at 301 nm.

### Solid Phase Peptide Synthesis

Employing a previously outlined approach,^31^ linear peptides were synthesized on a 0.05 mmol scale utilizing a PS3 peptide synthesizer (Gyros Protein Technologies). Preloaded 2-chlorotrityl chloride (CTC) resin, as detailed in resin loading procedures, was employed. Fmoc deprotection was carried out using a 20% piperidine/DMF solution (2 x 5 minutes). Dicyclohexylcarbodiimide (DIC), Oxyma Pure, and Fmoc-protected amino acids (each 6 equivalents) were utilized in a 1-hour coupling process. The coupling and deprotection steps were reiterated until the desired linear peptide sequence was achieved.

### Resin Cleavage

Following a previously described method,^31^ the resin obtained from solid phase peptide synthesis (SPPS) was exposed to DMF for 15 minutes to facilitate resin swelling within a 5 mL fritted polypropylene syringe (Torviq). Subsequently, the DMF was drained, and the resin was treated with a 20% piperidine/DMF solution for 15 minutes. Afterward, the resin was washed with DMF (3 x 2 mL) and DCM (3 x 2 mL), followed by drying. To confirm successful Fmoc deprotection, a Kaiser ninhydrin test was conducted. The deprotected peptide was further treated with 25% HFIP/DCM for 30 minutes. The resultant solution was concentrated under vacuum, and the residue was resuspended in 50% H2O/MeCN, frozen, and lyophilized. The crude compound obtained was used without additional purification.

### Peptide Cyclization

Adhering to a previously established procedure,^31^ crude peptides (∼0.05 mmol), benzotriazolyloxy-tris[pyrrolidino]-phosphonium hexafluorophosphate (PyBop, 3 equivalents), DIPEA (6 equivalents), and DMF (1.25 mM) were subjected to overnight agitation (16-24 hours) and subsequently concentrated under vacuum. To the concentrated crude residue, 10 mL of 50% H2O/ acetonitrile (MeCN) was added, vortexed, and centrifuged to isolate the precipitate and decant the supernatant. The resulting precipitate underwent additional washing with another 10 mL of 50% H2O/MeCN, followed by freezing and lyophilization. In case the residue did not precipitate, prep HPLC was employed to obtain the cyclized product.

### Global Deprotection

Following a previously described protocol,^31^ the crude cyclized product underwent treatment with trifluoroacetic acid (TFA): DCM: triisopropylsilyl (TIPS) (50: 45: 5, 20 mM) for 2 hours. The volatiles were removed using air, and the peptides were precipitated with 2 mL of Methyl tert-butyl ether (TBME). The precipitate was vortexed, centrifuged, washed with additional TBME, dissolved in H2O: MeCN, frozen, and lyophilized.

### HPLC

Following a previously documented procedure,^31^ High-performance liquid chromatography (HPLC) analysis and purification were conducted using an Agilent Technologies 1260 Infinity II preparative HPLC system. Purity analysis utilized a Luna C18 reverse phase column (5 μm, 150 x 4.6 mm, Phenomenex), while purification was carried out employing a Luna C18 reverse phase column (5 μm, 150 x 21.2 mm, Phenomenex). Eluent A consisted of H_2_O with 0.1% formic acid (FA), while Eluent B comprised MeCN with 0.1% FA. The gradient for purity analysis was as follows: (A:B, flow rate 1 mL/min): 95:5, 0 min; 95:5, 1 min; 5:95, 20 min; 5:95, 25 min; 95:5, 30 min. For purification, the gradient was adjusted: (A:B, flow rate 20 mL/min): 95:5, 0 min; 95:5, 1 min; 5:95, 20 min; 5:95, 25 min; 95:5, 30 min.

## Result and Discussion

### Cyclic Peptides Exhibited Significant Trophocidal Activity against *B. mandrillaris*

*B. mandrillaris* is a neglected infectious disease with a high mortality rate. As such, few studies for molecules that are effective against *B. mandrillaris* have been performed.^9,11,12,32,33^ Among the currently recommended drugs, only pentamidine has significant *in vitro* activity against *B. mandrillaris*, with a reported IC_50_ of between 9 and 18 μM.^12,34^ To date, no one has shown curative *in vivo* activity due to the lack of robust translational pathobiological models. Recently, nitroxoline was screened against *B. mandrillaris* trophozoites *in vitro* and was identified to have an IC_50_ of 4.8 μM.^12^ Since the publication of these results, nitroxoline was successfully repurposed to treat a case of BAE disease in California, US.^35^ The patient improved after nitroxoline administration and excitingly survived.^12,35^ However, kidney toxicity complications were observed. Given this dearth of effective, well-tolerated *B. mandrillaris* treatments, and the great utility that cyclic peptides have shown as treatments for other diseases, we screened 44 cyclic peptides from the SNaPP library at 16 μg/mL for trophocidal activity against *B. mandrillaris, Acanthamoeba castellanii*, and *Naegleria fowleri*. 8 hits for *B. mandrillaris*, 16 hits for *Acanthamoeba castellanii*, and one hit for *Naegleria fowleri* were identified and validated via dose-response, giving us a hit rate of 18%, 50%, and 2%, respectively (**Figure 2A** and **Table S1-S2**). Of the identified hits, the most promising were pNP-43 for *B. mandrillaris*, pNP-51 for *A. castellanii*, and pNP-12 for *N. fowleri*. We chose to go forward with pNP-43 given its strong initial potency (IC_50_ = 5 μM). Of the initial 8 *B. mandrillaris* hits, 6 were closely structurally related to pNP-43, providing further support for pursuing this scaffold (**Figure 2B**).

**Figure 2.**
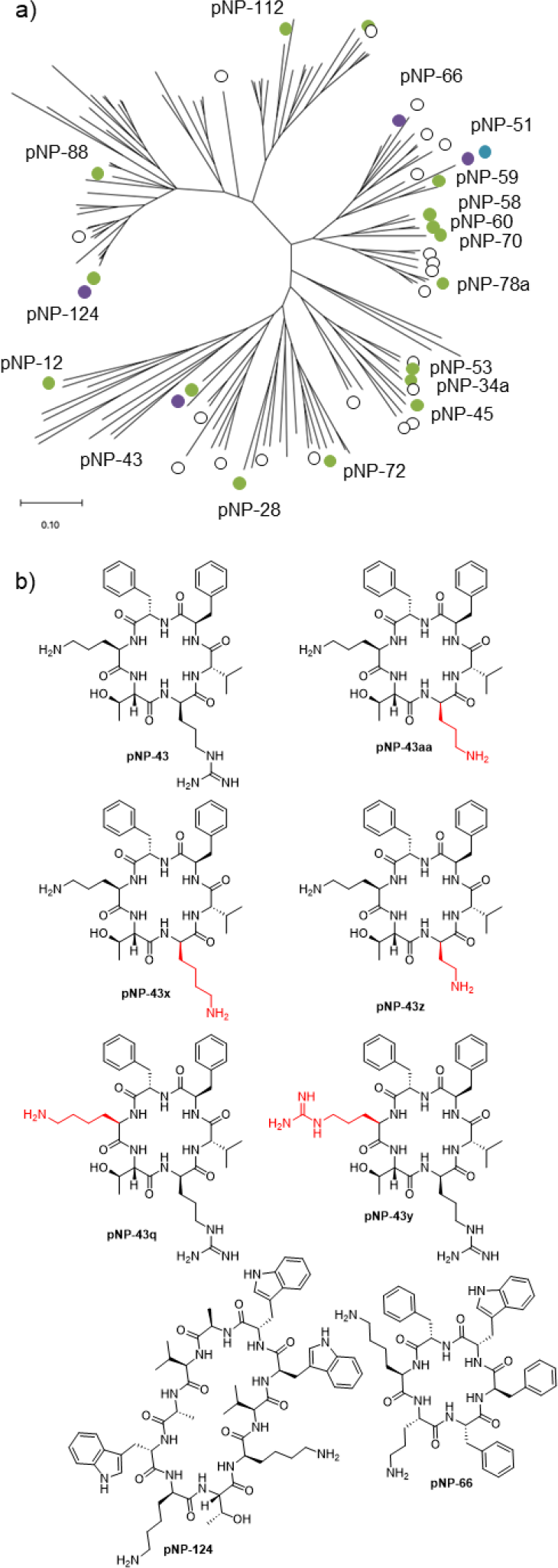
a) Tanimoto similarity tree of the previously predicted peptide library. All circles are compounds tested. Purple depicts hits against *Balamuthia mandrillaris*, green depicts peptides active against *Acanthamoeba castellanii*, and blue depicts cyclic peptides that inhibits *Naegleria fowleri*. All hits have their designated pNP number next to their respective circle. b) Structures of the top 8 initial hits against *Balamuthia mandrillaris*. Of the top 8, 6 are derivatives of pNP-43.

We next performed an alanine scan for each position of the cyclic hexapeptide to identify residues that are necessary for activity (**Table 1**). While all amino acids were found to contribute to the *B. mandrillaris* inhibitory activity, the D-orthinine (position 2 of pNP-43) and the threonine (position 3 of pNP-43) were found to not be essential (pNP43b and pNP43c, respectively). A linear version of pNP-43 (pNP-43g) was also tested, but it demonstrated a significant decrease in activity, confirming that the cyclic conformation is essential for activity. Following the alanine scan, further derivatization of pNP-43 was performed to identify more potent analogues. Modifying either of the phenylalanines (position 1 or 6 of pNP-43) with either a 4-flurophenylalanine (pNP-43m or pNP-43i) or 3,4-flurophenylalanine (pNP-43n) or pNP-43j) resulted in a decrease in activity (**Table S3**). However, when a 3-flurophenylalanine (pNP-43h or pNP-43l) was substituted at this position, the activity was maintained, suggesting that an electron withdrawing group is tolerated in the *meta* position, but not the *para* position. Tyrosine substitution at the one position (pNP-43k) maintained activity but was not tolerated at the six position (pNP-43o).

**Table 2.** Structures, potency, and toxicity analyses for the alanine scan and top compounds.

| Name | Structure | *B. mandrillaris*<br>IC<sub>50</sub> (μM) | A549<br>IC<sub>50</sub> (μM) | SI | Hemolysis<br>at 100 μM |
| --- | --- | --- | --- | --- | --- |
| pNP-43 | c[Phe-D-Orn-Thr-D-Arg-Val-D-Phe] | 4.6 ± 0.2 | >100 | >22 | <10% |
| pNP-43a | c[Ala-D-Orn-Thr-D-Arg-Val-D-Phe] | > 20 | ND | ND | ND |
| pNP-43b | c[Phe-D-Ala-Thr-D-Arg-Val-D-Phe] | 11.9 ± 1.8 | 48 | 4 | <10% |
| pNP-43c | c[Phe-D-Orn-Ala-D-Arg-Val-D-Phe] | 8.5 ± 0.7 | >100 | >12 | <10% |
| pNP-43d | c[Phe-D-Orn-Thr-D-Ala-Val-D-Phe] | > 20 | ND | ND | ND |
| pNP-43e | c[Phe-D-Orn-Thr-D-Arg-Ala-D-Phe] | > 20 | ND | ND | ND |
| pNP-43f | c[Phe-D-Orn-Thr-D-Arg-Val-D-Ala] | > 20 | ND | ND | ND |
| pNP-43g | H-Val-D-Phe-Phe-D-Orn-Thr-D-Arg-OH | > 20 | ND | ND | <10% |
| pNP-43h | c[Phe(3F)-D-Orn-Thr-D-Arg-Val-D-Phe] | 4.7 ± 1.0 | >100 | >21 | ND |
| pNP-43p | c[Phe-D-Lys-Thr-D-His-Val-D-Phe] | 2.4 ± 0.1 | ND | ND | <10% |
| pNP-43x | c[Phe-D-Orn-Thr-D-Lys-Val-D-Phe] | 3.4 ± 0.6 | ND | ND | <10% |
| pNP-43s | c[Phe-D-Lys-Thr-D-Lys-Val-D-Phe] | 3.9 ± 0.5 | 98 | 25 | <10% |
| pNP-43q | c[Phe-D-Lys-Thr-D-Arg-Val-D-Phe] | 4.1 ± 0.6 | 71 | 17 | <10% |
| pNP-43l | c[Phe-D-Orn-Thr-D-Arg-Val-D-Phe(3F)] | 4.2 ± 0.8 | 64 | 15 | <10% |
| pNP-43y | c[Phe-D-Arg-Thr-D-Arg-Val-D-Phe] | 4.4 ± 0.5 | ND | ND | <10% |

We then explored varying the chain length of nucleophilic amino acids at position 2 and 4. First, D-arginine at position 4 was exchanged for an amino acid with a shorter chain length, such as D-lysine (pNP-43x), D-orninthine (pNP-43aa), and D-diaminobutyric acid (pNP-43z). While D-lysine (pNP-43x) was well tolerated, the other modifications were not, suggesting that a side chain with 4 or more carbons at position 4 is important for activity. When the D-ornithine at position 6 was modified with D-lysine (pNP-43q) and D-arginine (pNP-43y), resulting in a longer chain length, the inhibitory activity was maintained. However, shortening the chain with a D-diaminopropionic acid (pNP-43t) and D-diaminobutyric acid (pNP-43u) resulted in decreased activity. This suggests that a chain length of at least 3 carbons between the alpha-carbon and the basic residue is important for inhibitory activity. Interestingly, these results are similar to those previously found in the study of cationic quarternary ammounium compounds and alkylphosphocholines.^36^ While these molecules are quite structurally different from the cyclic peptides studied here, they also showed increased activity with increasing alkyl chain length. This points to the activity of these molecules being due, at least in part, to their ability to interact with the protist plasma membrane. This is consistent with one of the major mechanisms of antimicrobial peptides, interaction with the cell wall through hydrophobic or electrostatic interactions, leading to membrane damage or lysis, cytoplasmic leakage, and cell death.^37–40^ Future studies will investigate the mechanisms of action of the *B. mandrillaris* inhibitory peptides in more detail.

When position 4 is substituted with a D-histidine (CN-1-117, pNP-43v), activity decreases. However, when this modification is combined with position 2 being changed to a D-lysine (CS-1-50, pNP-43p) activity increases. For this reason, we explored additional substitutions of position 4 while keeping position 2 a D-lysine. Changing position 4 to D-lysine (CN-1-121, pNP-43s) maintains activity while D-tryptophan (CN-1-115, pNP-43r) decreases activity. This further confirms that a chain length greater than 4 is needed at position 4 and suggests that a locked conformation is not tolerated at that position. Given this information, a structure activity relationship was determined (**Figure 3A**). Excitingly, 8 compounds were identified with IC_50_s of less than 5 μM (pNP-43p, pNP-43x, pNP-43s, pNP-43q, pNP-43l, pNP-43y, pNP-43g, and pNP-43u). The top hit, pNP-43p, was chosen to directly compare the *in vitro* performance against pentamidine. pNP-43p showed more potent efficacy (∼with the IC_50_ of 2.43 μM, compared to the IC_50_ of 6.59 μM for pentamidine (**Figure 3B**).

**Figure 3.**
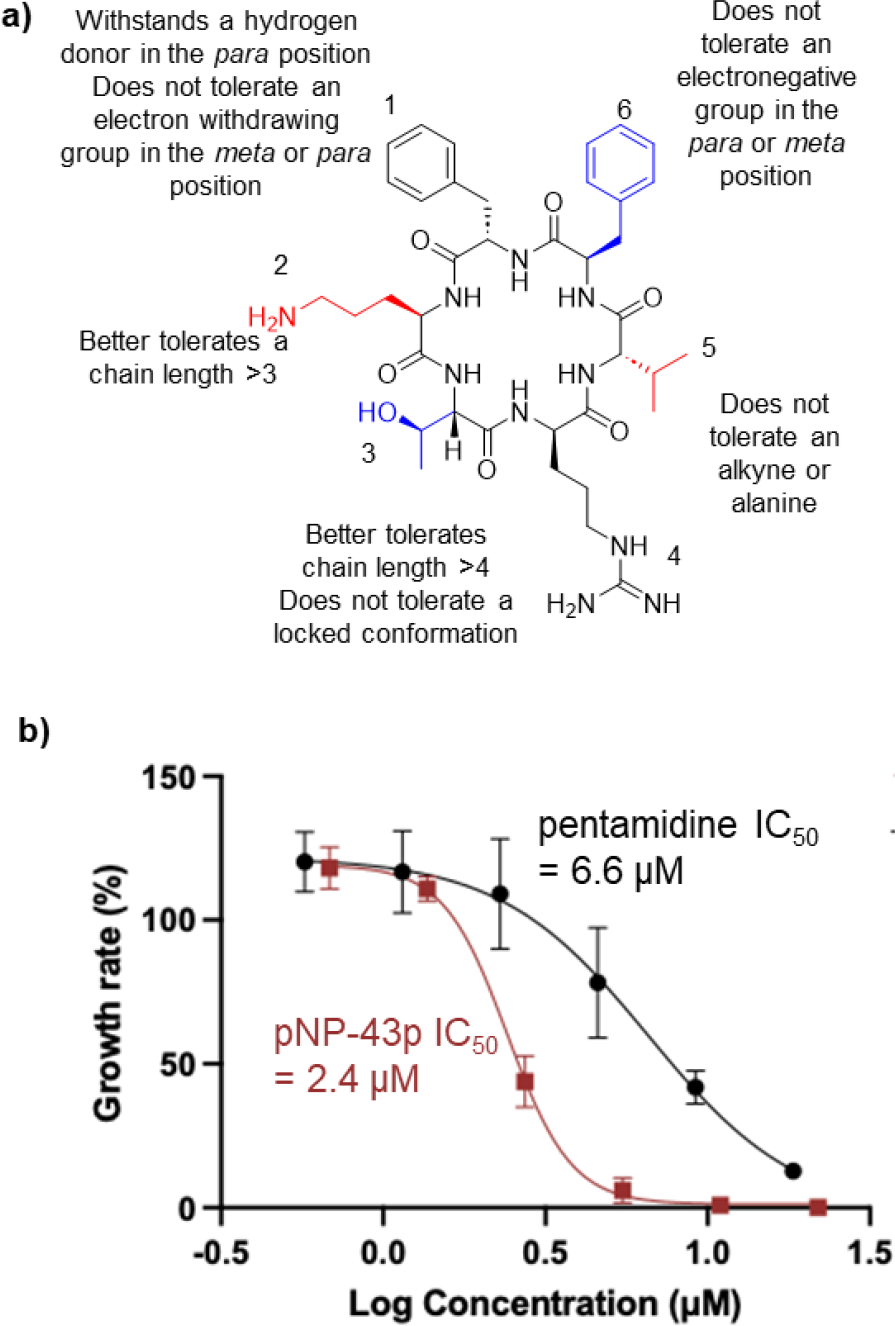
a) Structure activity relationship of pNP-43. B) Dose response curve for the most active derivative (pNP-43p) and pentamidine in the *B. mandrillaris* activity assay.

We next decided to screen for the ability of these molecules to inhibit the growth of other FLA. Interestingly, neither our initial hit (pNP-43) nor the derivatives tested showed strong activity (IC_50_ < 10 μM, **Table S2**). The selectivity of these cyclic peptides for *B. mandrillaris* over *Acanthamoeba* and *Naegleria* is quite interesting and points to a unique mechanism of action. It is possible that this selectivity may be due to differences in the composition of plasma membranes, given that many cyclic peptides act at the plasma membrane. However, additional studies are needed to confirm this.

### Cyclic Peptides Exhibited Minimal Toxicity against Human Cells

Many of the currently used medicines for *B. mandrillaris* have severe toxicities. For example, a 57-year-old cutaneous *B. mandrillaris* infected patient died with severe liver damage potentially due to complications of the diminazene aceturate treatment they received.^41^ Another patient with *B. mandrillaris* GAE was treated for more than 2 months with the recommended multiple-drug regimen (flucytosine, pentamidine, fluconazole, sulfadiazine, azithromycin, miltefosine, and albendazole) and developed adverse effects, including acute kidney injury and myelosuppression.^35,42^ These examples indicate that new treatments with lower toxicities are needed for *B. mandrillaris*. Cyclic peptide natural products are crucial sources of drug candidates because of their wide range of activities as well as their oftentimes low toxicities. Given the promising activity of several cyclic peptides against *B. mandrillaris*, we chose to evaluate the most promising compounds (IC_50_ < 5 μM) for their selectivity. Specifically, we chose to examine their cytotoxicity against a human cell line (A549) and their hemolytic activity (**Table 1**). Gratifyingly, all hit compounds tested showed minimal toxicity to A549 lung carcinoma cells (IC_50_ ≥ 64 μM). The selectivity index (SI = A549 IC_50_/*B. mandrillaris* IC_50_) was determined for each compound, with the lead compounds having SI ≥ 15. Additionally, none of the compounds caused hemolysis at the highest concentration tested (100 μM). Overall, this suggests that the cyclic peptides are likely to have minimal toxicity to humans and thus are promising leads going forward.

In conclusion, we have screened the SNaPP library to identify inhibitors of *B. mandrillaris*. The initial screen revealed 8 cyclic peptides with an IC_50_ less 16 μg/mL. Derivatives were then synthesized, enabling the development of a robust structural activity relationship. From this work, multiple derivatives were identified to have an IC_50_ of less than 5 μM and an IC_90_ of less than 10 μM, with minimal to no toxicity to mammalian cells. Given that pentamidine, one of the current gold standards of *B. mandrillaris* treatment, has an IC_50_ of 6.6 μM, these cyclic peptides show great promise as leads for this challenging to treat disease. Future studies will focus on mechanism-of-action and in vivo testing. Overall, this is the first study to demonstrate the effectiveness of peptides against *B. mandrillaris*, providing a new direction of anti-*B. mandrillaris* drug development.

## Author Contributions

S.N., C.L., E.I.P., and C.A.R. designed the experiments. S.N., G.C., and C.N. synthesized the cyclic peptides. C.L. performed amoebicidal assays and the cytotoxicity assays. S.N. performed the hemolysis assays. S.N., C.L., E.I.P., and C.A.R. writing, review, and editing. All authors contributed to the article and approved the submitted version.

