## Supplemental Files for "Discovery of cyclic peptide natural product inhibitors of *Balamuthia mandrillaris*"

**Supplementary Information**

### Supplementary Information

#### Table S1. Screening data of SNaPP peptides with *B. mandrillaris*

| Name | Structure | $IC_{50}$ (μg/mL) | $IC_{90}$ (μg/mL) |
|---|---|---|---|
| pNP-12 | c[Val-Phe-D-Arg-D-Ser] | > 16.0 | > 16.0 |
| pNP-19 | c[Phe-D-Phe-Val-D-Arg-D-Thr] | > 16.0 | > 16.0 |
| pNP-23 | c[Thr-Gly-Phe-D-Phe-End] | > 16.0 | > 16.0 |
| pNP-28 | c[Thr-D-Leu-D-Phe-End-D-Orn] | > 16.0 | > 16.0 |
| pNP-30 | c[Phe-D-Phe-Val-D-Arg-Thr-D-Thr] | > 16.0 | > 16.0 |
| pNP-34 | c[Gly-Phe-D-Val-D-Arg-Lys-D-Phe] | > 16.0 | > 16.0 |
| pNP-34a | c[Phe-D-Val-D-Orn-Gln-D-Phe-Gly] | > 16.0 | > 16.0 |
| pNP-35 | c[Gly-Leu-D-Val-D-Orn-Lys-D-Phe] | > 16.0 | > 16.0 |
| pNP-36 | c[Val-d-Val-Asn-D-Thr-Arg-D-Phe] | > 16.0 | > 16.0 |
| pNP-37 | c[Gly-Phe-D-Val-D-Orn-Lys-D-Phe] | > 16.0 | > 16.0 |
| pNP-43 | c[Val-D-Phe-Phe-D-Orn-Thr-D-Arg] | 3.96 | 4.1 |
| pNP-43x | c[Phe-D-Orn-Thr-D-Lys-Val-D-Phe] | 2.51 | 3.58 |
| pNP-43y | c[Phe-D-Arg-Thr-D-Arg-Val-D-Phe] | 3.54 | 4.81 |
| pNP-43z | c[Phe-D-Orn-Thr-D-Dab-Val-D-Phe] | 7.11 | 7.97 |
| pNP-43aa | c[Phe-D-Orn-Thr-D-Orn-Val-D-Phe] | 8.7 | 10.4 |
| pNP-45 | c[Gly-Arg-D-Val-D-Orn-Lys-D-Trp] | > 16.0 | > 16.0 |
| pNP-48 | c[Phe-D-Val-Asn-D-Orn-Arg-D-Trp] | > 16.0 | > 16.0 |
| pNP-51 | c[Phe-D-Orn-Orn-D-Orn-Phe-D-Phe] | 11.2 | > 16.0 |
| pNP-53 | c[Gly-Phe-D-Val-D-Orn-Lys-D-Trp] | > 16.0 | > 16.0 |
| pNP-57 | c[Trp-Ala-D-Asp-Orn-D-Lys-Val-D-Phe] | > 16.0 | > 16.0 |
| pNP-58 | c[Trp-Ala-D-Asp-Orn-D-Lys-Val-D-Trp] | > 16.0 | > 16.0 |
| pNp-59 | c[Phe-Thr-D-Lys-Lys-D-Orn-Phe-D-Phe] | > 16.0 | > 16.0 |
| pNP-60 | c[Trp-Ala-D-Orn-Lys-D-Lys-Val-D-Phe] | > 16.0 | > 16.0 |
| pNP-61 | c[Phe-Thr-D-Orn-Arg-D-Orn-Thr-D-Phe] | > 16.0 | < 0.500 |
| pNP-62 | c[Trp-Ala-D-Orn-Orn-D-Lys-Val-D-Phe] | > 16.0 | > 16.0 |
| pNP-63 | c[Trp-Ser-D-Lys-Lys-D-Lys-Phe-D-Trp] | > 16.0 | > 16.0 |
| pNP-64 | c[Phe-Ser-D-Lys-Orn-D-Orn-Trp-D-Phe] | > 16.0 | > 16.0 |
| pNP-66 | c[Trp-Phe-D-Lys-Orn-D-Orn-Phe-D-Phe] | 3.67 | 9.92 |
| pNP-69 | c[Phe-Thr-D-Orn-Val-D-Orn-Phe-D-Phe] | > 16.0 | > 16.0 |
| pNP-70 | c[Trp-Ser-D-Orn-Orn-D-Lys-Thr-D-Trp] | > 16.0 | > 16.0 |
| pNP-72 | c[D-Val-Thr-Phe-Gly-D-Arg-Ser-D-Thr] | NT | NT |
| pNP-76 | c[Phe-Thr-D-Lys-Val-D-Orn-Thr-D-Phe] | > 16.0 | > 16.0 |
| pNP-78a | c[Phe-Thr-Asp-D-Lys-Thr-D-Phe] | > 16.0 | > 16.0 |
| pNP-79* | c[Phe-Thr-D-Lys-Orn-D-Orn-Thr-D-Phe] | > 16.0 | > 16.0 |
| pNP-81 | c[Ala-D-Val-Val-Trp-Orn-D-Tyr-Trp-Ser] | > 16.0 | > 16.0 |
| pNP-88 | c[Trp-D-Ala-D-Ala-Trp-D-Orn-Thr-D-Orn-D-Phe] | > 16.0 | > 16.0 |
| pNP-96 | c[Val-D-Val-D-Val-D-Val-Val-Lys-D-Asn-Orn-D-Asn] | > 16.0 | > 16.0 |
| pNP-112 | c[Val-D-Val-D-Thr-Val-D-Val-Thr-Orn-D-Orn-Phe-D-Asn] | > 16.0 | > 16.0 |
| pNP-115 | c[Trp-D-Ala-D-Val-D-Ala-Phe-D-Orn-Thr-D-Orn-Val-D-Trp] | > 16.0 | > 16.0 |
| pNP-124 | c[Trp-D-Ala-D-Val-D-Ala-Trp-D-Lys-Thr-D-Lys-Val-D-Trp] | 7.82 | 13.7 |
| WP_049978124.1 | c[Arg-D-Orn-Thr-D-Leu-D-Phe] | > 16.0 | > 16.0 |
| WP_099869982.1 | c[Phe-D-Phe-D-Arg-Thr-Gly] | > 16.0 | > 16.0 |
| ELQ80643.1 | c[Phe-D-Phe-D-Orn-Lys-Gly] | > 16.0 | > 16.0 |
| WP_086708205.1 | c[Trp-D-Orn-Thr-D-Orn-Trp-D-Ala-D-Val-D-Ala] | > 16.0 | > 16.0 |

Supplementary Information

**Table S2. Screening data of SNaPP peptides with *N. fowleri* and *A.castellanii*.**

| Name | Structure | *Naegleria fowleri* | | *Acanthamoeba castellanii* | |
|---|---|---|---|---|---|
| | | $IC_{50}$ (µg/mL) | $IC_{90}$ (µg/mL) | $IC_{50}$ (µg/mL) | $IC_{90}$ (µg/mL) |
| pNP-12 | c[Val-Phe-D-Arg-D-Ser] | > 16.0 | > 16.0 | 5.27 | > 16.0 |
| pNP-19 | c[Phe-D-Phe-Val-D-Arg-D-Thr] | > 16.0 | > 16.0 | > 16.0 | > 16.0 |
| pNP-23 | c[Thr-Gly-Phe-D-Phe-End] | NT | NT | NT | NT |
| pNP-28 | c[Thr-D-Leu-D-Phe-End-D-Orn] | > 16.0 | > 16.0 | 12.1 | > 16.0 |
| pNP-30 | c[Phe-D-Phe-Val-D-Arg-Thr-D-Thr] | > 16.0 | > 16.0 | > 16.0 | > 16.0 |
| pNP-34 | c[Gly-Phe-D-Val-D-Arg-Lys-D-Phe] | > 16.0 | > 16.0 | > 16.0 | > 16.0 |
| pNP-34a | c[Phe-D-Val-D-Orn-Gln-D-Phe-Gly] | > 16.0 | > 16.0 | 15.8 | > 16.0 |
| pNP-35 | c[Gly-Leu-D-Val-D-Orn-Lys-D-Phe] | > 16.0 | > 16.0 | > 16.0 | > 16.0 |
| pNP-36 | c[Val-d-Val-Asn-D-Thr-Arg-D-Phe] | NT | NT | > 16.0 | >16.0 |
| pNP-37 | c[Gly-Phe-D-Val-D-Orn-Lys-D-Phe] | NT | NT | NT | NT |
| pNP-43 | c[Val-D-Phe-Phe-D-Orn-Thr-D-Arg] | > 16.0 | > 16.0 | 13.1 | > 16.0 |
| pNP-43x | c[Phe-D-Orn-Thr-D-Lys-Val-D-Phe] | > 16.0 | > 16.0 | > 16.0 | > 16.0 |
| pNP-43y | c[Phe-D-Arg-Thr-D-Arg-Val-D-Phe] | > 16.0 | > 16.0 | > 16.0 | >16.0 |
| pNP-43z | c[Phe-D-Orn-Thr-D-Dab-Val-D-Phe] | > 16.0 | > 16.0 | 12.6 | > 16.0 |
| pNP-43aa | c[Phe-D-Orn-Thr-D-Orn-Val-D-Phe] | > 16.0 | > 16.0 | > 16.0 | > 16.0 |
| pNP-45 | c[Gly-Arg-D-Val-D-Orn-Lys-D-Trp] | > 16.0 | > 16.0 | 10.5 | > 16.0 |
| pNP-48 | c[Phe-D-Val-Asn-D-Orn-Arg-D-Trp] | > 16.0 | > 16.0 | > 16.0 | > 16.0 |
| pNP-51 | c[Phe-D-Orn-Orn-D-Orn-Phe-D-Phe] | 5.98 | > 16.0 | > 16.0 | > 16.0 |
| pNP-53 | c[Gly-Phe-D-Val-D-Orn-Lys-D-Trp] | > 16.0 | > 16.0 | 12.1 | > 16.0 |
| pNP-57 | c[Trp-Ala-D-Asp-Orn-D-Lys-Val-D-Phe] | > 16.0 | > 16.0 | > 16.0 | > 16.0 |
| pNP-58 | c[Trp-Ala-D-Asp-Orn-D-Lys-Val-D-Trp] | > 16.0 | > 16.0 | 14.8 | > 16.0 |
| pNp-59 | c[Phe-Thr-D-Lys-Lys-D-Orn-Phe-D-Phe] | > 16.0 | > 16.0 | 12.9 | > 16.0 |
| pNP-60 | c[Trp-Ala-D-Orn-Lys-D-Lys-Val-D-Phe] | > 16.0 | > 16.0 | 13.6 | > 16.0 |
| pNP-61 | c[Phe-Thr-D-Orn-Arg-D-Orn-Thr-D-Phe] | > 16.0 | >16.0 | > 16.0 | >16.0 |
| pNP-62 | c[Trp-Ala-D-Orn-Orn-D-Lys-Val-D-Phe] | NT | NT | NT | NT |
| pNP-63 | c[Trp-Ser-D-Lys-Lys-D-Lys-Phe-D-Trp] | > 16.0 | > 16.0 | > 16.0 | > 16.0 |
| pNP-64 | c[Phe-Ser-D-Lys-Orn-D-Orn-Trp-D-Phe] | > 16.0 | > 16.0 | > 16.0 | > 16.0 |
| pNP-66 | c[Trp-Phe-D-Lys-Orn-D-Orn-Phe-D-Phe] | > 16.0 | > 16.0 | > 16.0 | > 16.0 |
| pNP-69 | c[Phe-Thr-D-Orn-Val-D-Orn-Phe-D-Phe] | > 16.0 | > 16.0 | > 16.0 | > 16.0 |
| pNP-70 | c[Trp-Ser-D-Orn-Orn-D-Lys-Thr-D-Trp] | > 16.0 | > 16.0 | > 16.0 | > 16.0 |
| pNP-72 | c[D-Val-Thr-Phe-Gly-D-Arg-Ser-D-Thr] | > 16.0 | > 16.0 | > 16.0 | > 16.0 |
| pNP-76 | c[Phe-Thr-D-Lys-Val-D-Orn-Thr-D-Phe] | NT | NT | NT | NT |
| pNP-78a | c[Phe-Thr-Asp-D-Lys-Thr-D-Phe] | > 16.0 | > 16.0 | 14.2 | > 16.0 |
| pNP-79* | c[Phe-Thr-D-Lys-Orn-D-Orn-Thr-D-Phe] | > 16.0 | > 16.0 | > 16.0 | > 16.0 |
| pNP-81 | c[Ala-D-Val-Val-Trp-Orn-D-Tyr-Trp-Ser] | > 16.0 | > 16.0 | > 16.0 | > 16.0 |
| pNP-88 | c[Trp-D-Ala-D-Ala-Trp-D-Orn-Thr-D-Orn-D-Phe] | > 16.0 | > 16.0 | 11.6 | > 16.0 |
| pNP-96 | c[Val-D-Val-D-Val-D-Val-Val-Lys-D-Asn-Orn-D-Asn] | NT | NT | NT | NT |
| pNP-112 | c[Val-D-Val-D-Thr-Val-D-Val-Thr-Orn-D-Orn-Phe-D-Asn] | > 16.0 | > 16.0 | 11.2 | > 16.0 |
| pNP-115 | c[Trp-D-Ala-D-Val-D-Ala-Phe-D-Orn-Thr-D-Orn-Val-D-Trp] | > 16.0 | > 16.0 | > 16.0 | > 16.0 |
| pNP-124 | c[Trp-D-Ala-D-Val-D-Ala-Trp-D-Lys-Thr-D-Lys-Val-D-Trp] | > 16.0 | > 16.0 | 15.4 | > 16.0 |
| WP_049978124.1 | c[Arg-D-Orn-Thr-D-Leu-D-Phe] | > 16.0 | > 16.0 | > 16.0 | > 16.0 |
| WP_099869982.1 | c[Phe-D-Phe-D-Arg-Thr-Gly] | > 16.0 | > 16.0 | > 16.0 | > 16.0 |
| ELQ80643.1 | c[Phe-D-Phe-D-Orn-Lys-Gly] | > 16.0 | > 16.0 | > 16.0 | > 16.0 |
| WP_086708205.1 | c[Trp-D-Orn-Thr-D-Orn-Trp-D-Ala-D-Val-D-Ala] | NT | NT | NT | NT |

**Supplementary Information**

**Table S3. IC50 of cyclic peptides against trophozoites of *B. mandrillaris in vitro.***

| Compound Name | *B. mandrillaris* Mean $IC_{50}$ (μM) ±SD | *B. mandrillaris* Mean $IC_{90}$ (μM) ±SD | A549 Mean $IC_{50}$ (μM) | Selectivity Index (SI) |
|---|---|---|---|---|
| **pNP-43 (MH-2-08) Derivatives** | | | | |
| pNP-43p (CS-1-50) | 2.43 ± 0.04 | 4.02 ± 0.84 | ND | ND |
| pNP-43x (CS-1-03) | 3.41 ± 0.58 | 4.89 ± 0.15 | ND | ND |
| pNP-43s (CN-1-121) | 3.89 ± 0.54 | 6.65 ± 1.00 | 97.80 | 25.14 |
| pNP-43q (CN-1-119) | 4.11 ± 0.56 | 5.66 ± 0.04 | 70.50 | 17.15 |
| pNP-43l (CN-1-123) | 4.22 ± 0.77 | 5.89 ± 0.45 | 64.40 | 15.26 |
| pNP-43y (CS-1-06) | 4.38 ± 0.54 | 5.75 ± 0.46 | ND | ND |
| pNP-43 (SN-3-186) | 4.62 ± 0.17 | 7.38 ± 1.12 | >100 | >21.65 |
| pNP-43h (CN-1-127) | 4.68 ± 0.99 | 7.64 ± 1.94 | >100 | >21.37 |
| pNP-43 (MH-2-08) | 5.03 ± 0.06 | 5.73 ± 0.11 | ND | ND |
| pNP-43k (CN -1-133) | 5.41 ± 0.99 | 8.85 ± 1.62 | >100 | >18.48 |
| pNP-43c (SN-3-176) | 8.50 ± 0.73 | 10.90 ± 0.28 | >100 | >11.76 |
| CN-1-129 | 9.09 ± 1.24 | 12.45 ± 1.77 | >100 | >11.00 |
| GC-1-012 | 9.42 ± 0.03 | 10.85 ± 0.07 | >100 | >10.62 |
| CS-1-04 | 10.04 ± 0.16 | 11.23 ± 0.22 | ND | ND |
| SN-3-174 (pNP-43b) | 11.85 ± 1.77 | 12.50 ± 1.20 | 47.70 | 4.03 |
| CN-1-117 | 12.05 ± 2.62 | 13.10 ± 1.98 | 55.80 | 4.63 |
| CS-1-02 | 12.03 ± 0.10 | 14.39 ± 0.20 | ND | ND |
| CN-1-125 | 11.95 ± 1.77 | 17.85 ± 2.90 | >100 | >8.37 |
| GC-1-009 | 13.05 ± 0.21 | > 20.0 | 71.60 | 5.49 |
| CN-1-131 | 14.40 ± 2.40 | > 20.0 | >100 | >6.94 |
| GC-1-010 | 16.35 ± 1.06 | > 20.0 | >100 | >6.12 |
| GC-1-011 | 18.00 ± 1.41 | > 20.0 | 79.80 | 4.43 |
| **CS-1-13 Derivatives** | | | | |
| CS-1-46 | 3.50 ± 0.05 | 5.08 ± 0.12 | ND | ND |
| CS-1-59 | 10.21 ± 0.54 | 12.18 ± 0.26 | ND | ND |
| SN-2-136 | 12.59 ± 1.52 | >20.0 | ND | ND |
| CS-1-47 | 13.45 ± 0.20 | >20.0 | ND | ND |
| SN-2-132 | 14.11 ± 0.00 | >20.0 | ND | ND |
| CS-1-13 | 13.78 ± 1.26 | >20.0 | ND | ND |
| CS-1-45 | 14.78 ± 0.79 | >20.0 | ND | ND |
| SN-2-52 | 19.57 ± 2.48 | >20.0 | ND | ND |

ND - data not determined.
